# The anticipated potential nuclear localization sequence of ‘*Candidatus* Phytoplasma mali’ SAP11-like protein is required for TCP binding but not for transport into the nucleus

**DOI:** 10.1101/2020.05.13.094045

**Authors:** Alisa Strohmayer, Timothy Schwarz, Mario Braun, Gabi Krczal, Kajohn Boonrod

**Affiliations:** RLP AgroScience GmbH, AlPlanta Institute for Plant Research, Breitenweg 71, 67435 Neustadt, Germany

## Abstract

The plant pathogen ‘*Candidatus* Phytoplasma mali’ (‘*Ca*. P. mali’) is the causing agent of apple proliferation that leads to heavy damage in apple production all over Europe. To identify and analyze effector proteins of plant pathogens is an important strategy in plant disease research. Here, we report that the SAP11-like protein of *‘Ca*. P. mali’ induces crinkled leaves and siliques and witches’ broom symptoms in transgenic *Arabidopsis thaliana* (*A. thaliana*) plants and binds to 6 members of class I and all members of class II TCP (TEOSINE BRANCHES/ CYCLOIDEA/PROLIFERATING CELL FACTOR) transcription factors of *A. thaliana* in yeast two-hybrid assays. Moreover, we demonstrate that the protein localizes actively into the plant nucleus without requiring the nuclear leader sequence (NLS). We also identified a 17 amino acid stretch previously predicted to be a nuclear leader sequence that is important for the binding of some of the TCPs and also responsible for the crinkled leaf and silique phenotype in transgenic *A. thaliana*.

## Introduction

The delivery of effector proteins and small molecules into the plant host is a common strategy of plant pathogens, including bacteria, fungi, oomycetes and nematodes, to enhance the hosts’ susceptibility and benefit their infectiousness (1). The function of these effectors reaches from suppression of the plant immune system to alteration of plant behavior and development (1). Thus, identifying targets of plant pathogen effectors and revealing plant-microbe interactions enable to better understand the infectious mechanisms and consequently to control phytoplasma diseases.

Phytoplasmas are plant pathogenic bacteria that are transmitted by insect vectors and reside in the phloem of their plant host. Phytoplasmas are the causative agent of numerous diseases in plants, including important food crops, leading to heavy damage to the host plant, considerable yield loss, and eventual death of the plant. Phytoplasma has been shown to secrete effector proteins that change plant development and increase phytoplasma fitness (2, 3). Especially the genome of Aster Yellows phytoplasma strain Witches Broom (AY-WB) (4) was mined thoroughly for potential effector proteins by identifying proteins with N-terminal signal peptide (SP) sequence that are secreted via the Sec-depended pathway (5). One of these secreted AY-WB proteins (SAP) is SAP11, a small effector protein which has been extensively investigated. SAP11 specifically targets the plant cell nucleus via a nuclear localization sequence (NLS) within the protein and plant importin α (5). Transgenic *A thaliana* lines expressing AY-WB SAP11 show severe symptoms including crinkled leaves, crinkled siliques, stunted growth and an increase in stem number (6). The biochemical analysis of transgenic plants expressing AY-WB_SAP11 shows that SAP11 binds and destabilizes CINCINNATA (CIN)-related TEOSINTE BRANCHED1, CYCLOIDEA, PROLIFERATING CELL FACTORS (TCP) transcription factors leading to a decrease of jasmonate (JA) production and an enhanced insect vector reproduction (7, 6). Similar changes in phenotype development as well as a reduction of JA production and enhancement of insect progeny could also be found in *A. thaliana* plants infected with AY-WB phytoplasma (7).

In ‘*Candidatus* Phytoplasma mali’ (‘*Ca.* P. mali’) strain AT, a putative pathogenic-related effector protein ATP_00189 (GenBank: CAP18376.1) was identified which shares 41% homology on amino acid level with AY-WB_SAP11 and was therefore called SAP11-like protein (8). ‘*Ca*.’ P. mali is the cause of apple proliferation (AP) causing symptoms such as witches’ broom, enlarged stipules, tasteless and dwarf fruits and thus leading to massive yield losses and economic damage in apple production. AY-WB_SAP11 and the AP_SAP11-like protein share a signal-peptide motif of the phytoplasma-specific sequence-variable mosaic (SVM) protein signal sequence (Pfam entry: PF12113), linking these proteins to a rapid evolution (9). Both AY-WB_SAP11 and AP_SAP11-like protein are found to be expressed in infected plants (8, 5).

SAP11-like protein of ‘*Ca.* P. mali’ STAA, a strain found in northern Italy, binds TCP transcription factors and the infection with ‘*Ca.* P. mali’ STAA leads to altered phytohormonal levels in apple trees, including changes in JA, salicylic acid and abscisic acid levels (10). Furthermore, a change in the odor of the apple tree leads to an enhanced attraction of the insect vector (11, 12). A change in aroma phenotype, caused by the alteration of volatile organic compounds (VOC) production of the plant, is also detected in transgenic *Nicotiana benthamiana* (*N. benthamiana*) plants that express the SAP11-like protein of ‘*Ca*. P. mali’ (13). The similarities of TCP-binding, the biochemical changes in transgenic plants and hydrophobic amino acid patterns despite their differences in amino acid sequence of AY-WB_SAP11 and AP_SAP11-like protein, leads to the assumption that these proteins may have a similar function during phytoplasma infection (10). Due to difference of the amino acid feature of the predicted NLS of AP_SAP11-like protein compared to AY-WB_SAP11, the exact function of this domain was not analyzed so far. Recently, Wang and co-worker (14) shown that the NLS deletion of SAP11 like effector of wheat blue dwarf phytoplasma, SWPI, did not change the localization’s pattern of the protein which is contrary with the result of AY-WB SAP11.

Thus, these controversial facts prompt us to investigate a stretch of the AP_SAP11-like protein which corresponds to the predicted NLS of AY-WB_Sap11 (5) in sequence alignment in regards to the localization and the binding of AtTCP transcription factors. Furthermore, we compare our findings with current knowledge on SAP11 effectors phytoplasma and discuss its possible functions.

## Results

### Sequence analysis of SAP11-like protein of ‘*Candidatus* Phytoplasma mali’ strain PM19

The gene of the SAP11-like protein (GenBank Accession number MK966431) studied in this work was isolated from ‘*Ca*. P. mali’ strain PM19 (15). Its gene product is a 14 kDa protein composed of 122 amino acids. The alignment results of AP_SAP11-like_PM19 show a shared identity of 40 % with AY-WB_SAP11 (Accession Number WP_011412651) and a high similarity to AP_SAP11-like proteins of other strains (Fig. 1a). Compared to STAA (10) and AT (16) strain, there are only one and two amino acids substituted respectively (Fig. 1a).

**Figure 1:**
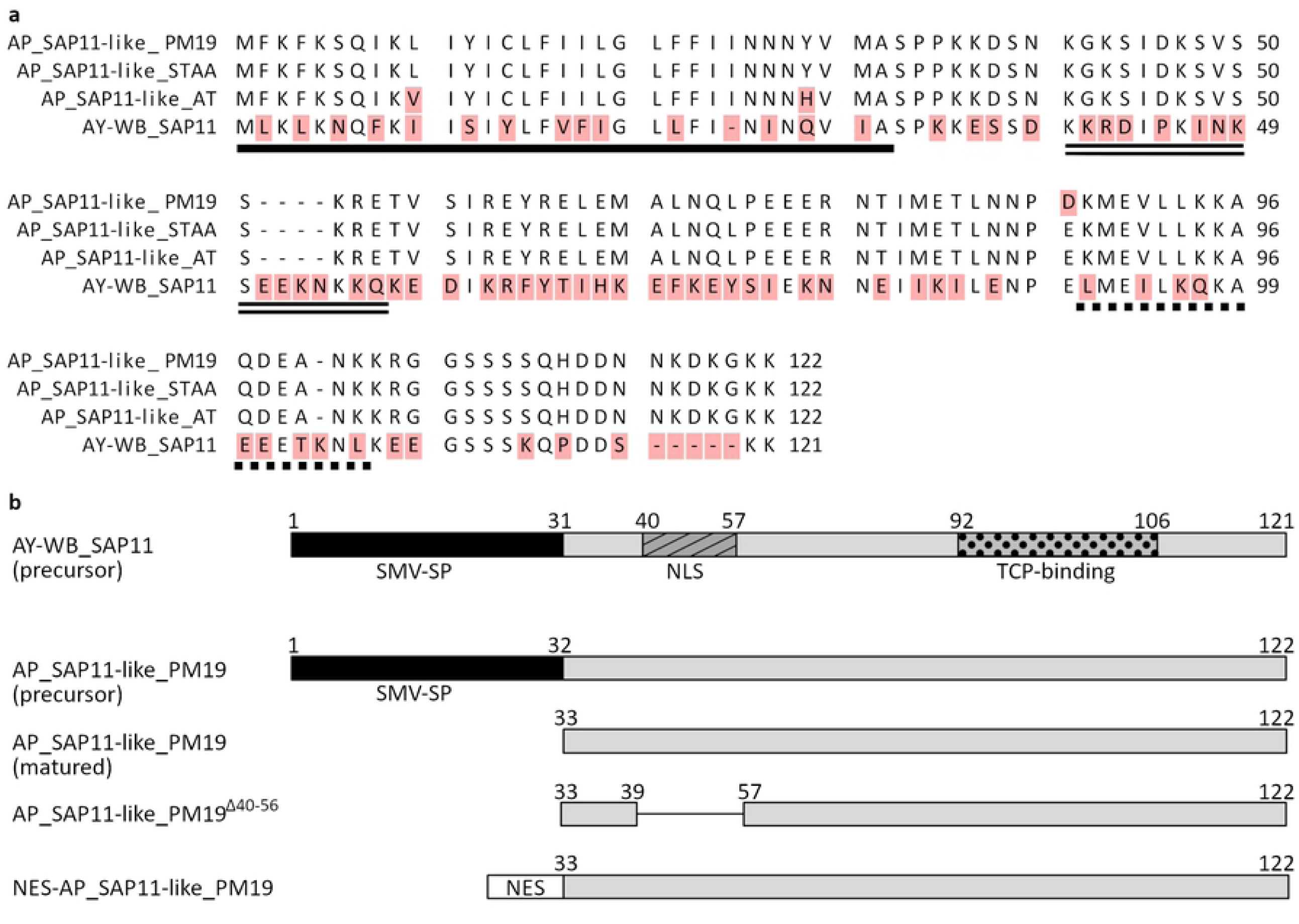
Schematic representation of the AP_SAP11-like_PM19 amino acid sequence used in this study. **a.** Multiple sequence alignment of AP_SAP11-like protein of strains PM19, STAA and AT and AY-WB_SAP11. Annotations: single black line represents sequence-variable mosaic protein signal peptide (SVM-SP), double black line represent nuclear leader sequence (NLS) and the dotted line represent TCP-binding area of AY-WB_SAP11. **b.** Schematic representation of deletions and additions to AP_SAP11-like _PM19comparison to AY-WB_SAP11. The black square represents the SVM-SP, the striated squares the NLS and the dotted squares the TCP-binding area. The deletion of amino acids 40 to 56 in AP_SAP11-like_PM19^Δ40-56^ is indicated by a line connecting the squares. In NES-AP_SAP11-like_PM19, a nuclear export sequence (NES) is fused to the N-terminus of the protein, indicated by a white square.

AY-WB_SAP11 and AP_SAP11-like_PM19 protein share a SVM signal sequence which is located in the N-terminal part of the proteins from amino acid 1 to 31 or 32 respectively and is important for the secretion of the proteins into the plant host cell via the phytoplasma Sec-dependent pathway (5, 8). The secretion of proteins via the Sec-dependent pathway is connected with cleavage of the signal peptide, resulting in the mature protein (17). Thus, a truncated version of AP_SAP11-like protein of strain PM19 starting with amino acid 32, called AP_SAP11-like_PM19, was used in all experiments (Fig. 1b), otherwise it will be indicated.

For AY-WB_SAP11 two more domains have been identified: a NLS (amino acids 40 to 57) that is required for nuclear localization, and a TCP-binding area (amino acids 92 to 106) that is part of a predicted coiled coil structure (amino acids 90 to 110) (5, 7). It was discussed that AP_SAP11-like protein and AY-WB_SAP11 share similar hydrophobicity motifs that could be responsible for similar functions of the proteins (10). However, it has not been experimentally determined so far.

### Amino acids 40 to 56 of AP_SAP11-like_PM19 are not necessary for nuclear localization in *N. benthamiana*

SAP11 of AY-WB phytoplasma was shown to localize in the plant cell nucleus (5). To elucidate whether SAP11-like protein of AP phytoplasma strain PM19 locates as well in the plant cell nucleus, the gene *AP_SAP11-like_PM19* was codon-optimized for expression in *A. thaliana*, synthesized (GeneCust, Ellange, Luxembourg) and fused to green fluorescence protein (*GFP*) resulting in *AP*_*SAP11-like-PM19-GFP*. To mark the nucleus of the plants cells, a bipartite nuclear leading sequence (biNLS) of the *Nicotiana tabacum* domains rearranged methyltransferase 1 (NtDRM1) (18) was fused to red fluorescence protein (RFP). The *biNLS-RFP* was transiently co-expressed with *AP*_*SAP11-like_PM19-GFP* in *N. benthamiana* using an *Agrobacterium*-mediated expression system. Two days after *Agrobacterium*-infiltration, protoplasts were isolated from the infiltrated leaves and analyzed by confocal microscopy using GFP and RFP filters. The results show that the AP_SAP11-like_PM19-GFP is mainly localized in the nucleus marked by biNLS-RFP (Fig.2a, left panel).

Amino acids 40 to 57 of AY-WB_SAP11 have been predicted to have a function as a bipartite nuclear leading sequence (5). Disruption of this NLS by deletion of amino acids 56 to 72 or amino acids 40 to 57 led to a distribution of the protein in cytoplasm and nucleus (5). The same effect was reported when only two lysines at position 55 and 56 of the protein were deleted (7). In contrary, deleting the predicted NLS of a SAP11-like effector from wheat blue dwarf phytoplasma, SWPI, does not affect the nuclear transport of the protein (14). We questioned whether AP_SAP11-like _PM19 was transported into the plant nucleus in a similar manner.

For AP_SAP11-like_PM19, we were not able to predict any NLS sequence using different software (19–21). Researches so far indicate that AY-WB_SAP11 and AP_SAP11-like protein have a similar function (10, 13, 6) and thus might also have a similar structure. We hypothesized, that a potential NLS of AP_SAP11-like protein might be in a similar region like the AY-WB_Sap11 NLS.

To elucidate, whether the area of AP_SAP11-like_PM19 corresponding to the predicted NLS of AY-WB SAP11 in sequence alignment (amino acids 40 to 56, Fig.1a) is necessary for the nuclear localization of the protein, we constructed a truncated version of *AP*_*SAP11-like_PM19* missing amino acids 40 to 56 (Fig.1b), which correspond to the predicted NLS of AY-WB_Sap11 sequence alignment (Fig. 1a), and fused the protein to GFP, resulting in *AP_SAP11-like_PM19*^*Δ40-56*^*-GFP*. The result of the protoplasts isolated from the infiltrated leaves shows that AP_SAP11-like_PM19^Δ40-56^-GFP still localizes in the plant cell nucleus (Fig.2a, right panel). Therefore, the result suggests that the amino acids 40-56-stretch of AP-SAP11-like_PM19 is not required for transporting the protein into the plant nucleus.

**Figure 2:**
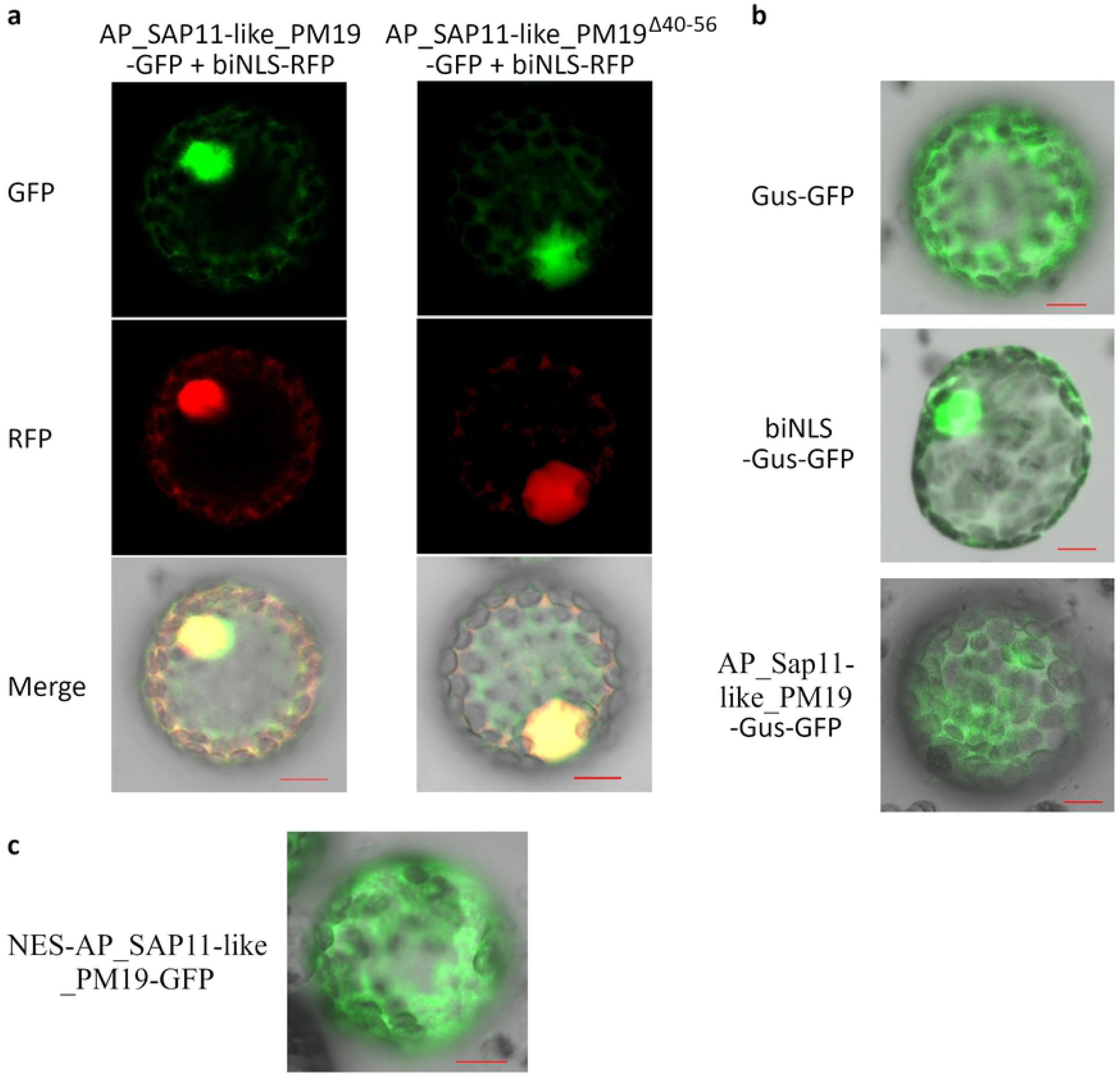
Localization studies of AP_SAP11-like_PM19 protein. Different constructs were fused to *GFP* or *RFP* and transiently expressed in *N. benthamiana*. Protoplasts of *Agrobacterium*-infiltrated leaves were analyzed by confocal microscopy using GFP and RFP filters. **a.** AP_SAP11-like_PM19 and AP_SAP11-like_PM19^Δ40-56^ localize mainly in the plant nucleus but also partly in the cytoplasm. Co-localization with biNLS-RFP is indicated by yellow coloring in the merge. **b.** Investigation of nuclear leading activity of AP_SAP11_PM19. Fusion of AP_SAP11-like_PM19 to Gus-GFP localizes mainly in the cytoplasm, whereas fusion with biNLS leads to a mainly nuclear localization. **c.** Fusion of AP_SAP11-like-PM19-GFP with a nuclear export sequence (NES) leads to cytoplasmic localization. Scale bar = 10 µm. Ten protoplasts expressing the proteins in questions were analyzed and all showed the same results.

### AP_SAP11-like protein fails to localize into nucleus when fused with GUS-GFP

Nuclear localization cannot only be achieved by active nuclear import, but also via passive transport. Even though it was long assumed that the maximum size for protein diffusion through nuclear pores is around 60 kDa, (22). Passive diffusion of the GFP (∼ 27 kDa) through nuclear pores is known and fusing a heterogeneous protein to GFP can lead to an unspecific signal in the nucleus (23). Thus, finding that AP_SAP11-like_PM19 and PM19^Δ40-56^-GFP localized in the nucleus could also be a false positive result caused by nuclear transportation of GFP. To prove this hypothesis, the AP_SAP11-like_PM19 was fused with β-Glucuronidase (Gus)-GFP. We used Gus-GFP fusion protein, since the expressed protein exclusively localizes in the cytoplasm when expressed alone (24). For a positive control, the biNLS of NtDRM1 (18) was fused to Gus-GFP. The result shows that Gus-GFP localized exclusively in cytoplasm while the biNLS-Gus-GFP is detected mainly in the nucleus (Fig.2b, upper and middle panel). As suspected, AP_SAP11-like_PM19 could not target the Gus-GFP into the nucleus. (Fig.2b, lower panel). These results suggest that the nuclear localization of AP_SAP11-like_PM19 is rather due to the size than the function of previously predicted NLS sequence (amino acids 40 to 56).

### AP_SAP11-like_PM19 induces crinkled leaves and siliques and witches’ broom symptoms in *Arabidopsis*

The expression of AY-WB_SAP11 in *A. thaliana* induces stem proliferation, leading to witches’ broom symptoms, and alteration of leaf and silique shape (6) while the expression of AP_SAP11-like protein of *Ca*. P. mali in *N. benthamiana* also leads to morphological changes with stunted growth and crinkled leaves (13).

To analyze the effects of AP_SAP11-like_PM19 protein in *A. thaliana*, we produced transgenic *Arabidopsis* plant lines stable expressing AP_SAP11-like_PM19 (Fig. 1b) under the control of the *Cauliflower mosaic virus* 35S promoter. The transgenic *A. thaliana* plant lines show smaller rosettes and a witches’ broom phenotype (Fig. 3a), crinkled leaves (Fig. 3b) and crinkled siliques (Fig. 3c)

**Figure 3:**
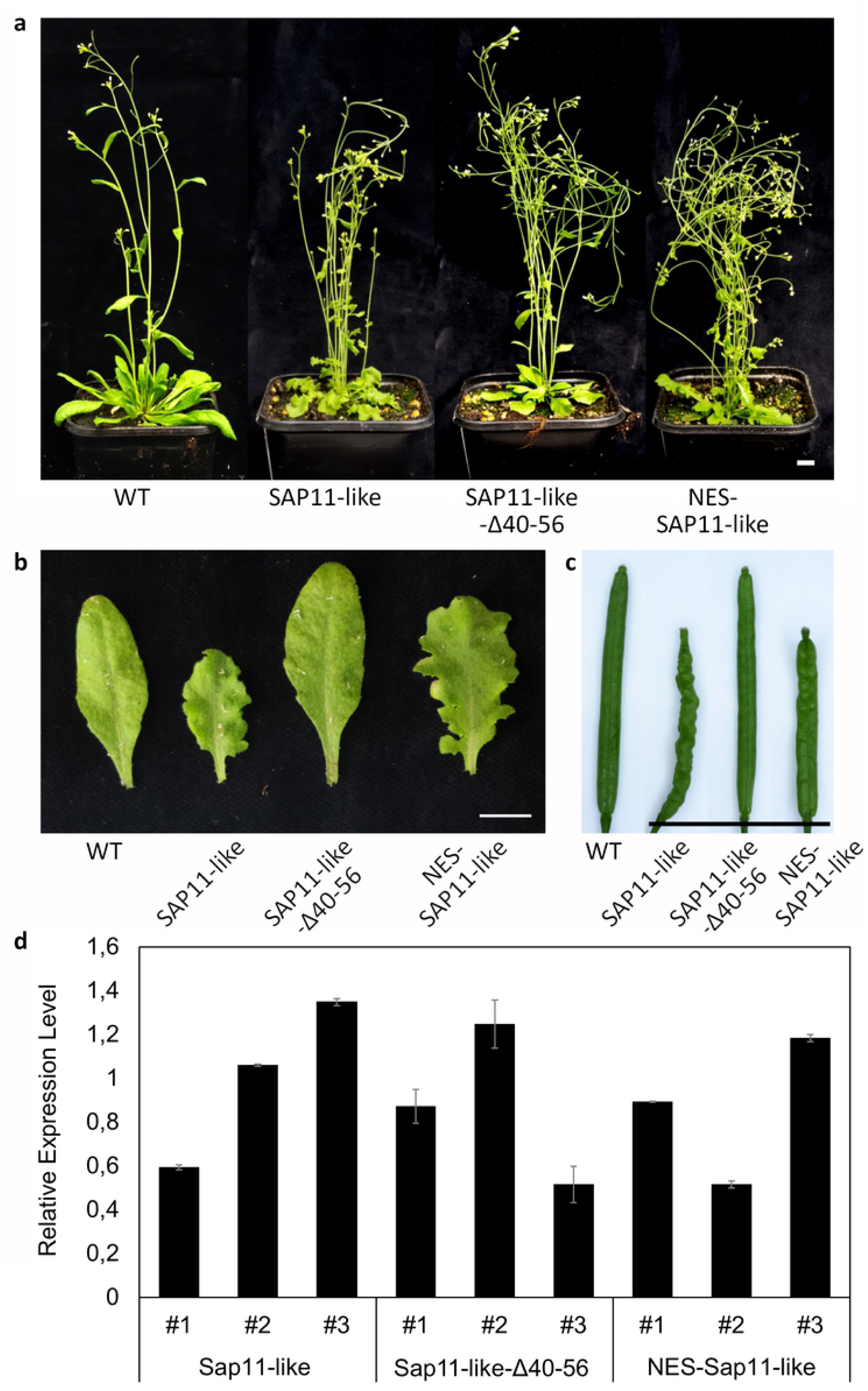
Amino acids 40 to 56 of AP_SAP11-like_PM19 protein are required for symptoms development in *A. thaliana*. **a.** Transgenic *A. thaliana* lines expressing AP_SAP11-like_PM19 (SAP11-like), AP_SAP11-like_PM19^Δ40-56^ (SAP11-like-Δ40-56) and NES-AP_SAP11-like_PM19 (NES-SAP11-like) under the control of the *Cauliflower mosaic virus* 35S promoter compared to *Arabidopsis* wild type Col-0 plant. All plants were grown for 8 weeks in long-day (16 h/8 h light/dark) conditions. For each of the three constructs, at least 10 lines were examined, all showing similar phenotypic characteristics. **b.** Leaves of transgenic plant lines shown in **a. c.** Siliques of transgenic lines shown in **a. d.** Relative expression levels of transgenes of transgenic lines compared to the geometric average of *glyceraldehyde-3-phosphate dehydrogenase* (*GAPDH*) and *protein phosphatase 2* (*PP2A*). #1 represents plants shown in **a.** respectively. All values show a P-value lower than 0.05 in ANOVA. Scale bars = 1 cm.

### Amino acids 40 to 56 of AP_ SAP11-like protein are important for symptom development

It was shown that transgenic *A. thaliana* lines expressing AY-WB_SAP11 missing the N-terminal region including the NLS or an AY-WB_SAP11 mutant lacking two lysines at position 55 and 56 loose the crinkled leave symptoms (7). However, it was not finally determined, whether the missing phenotype is caused by the lacking transport of the protein into the nucleus or by the mutation of the protein itself (Sugio et al. 2014).To reveal the role of the 40 to 56 amino acid region of AP_SAP11-like_PM19, we produced two transgenic *Arabidopsis* lines; (1) AP_SAP11-like_PM19^Δ40-56^ (Fig. 1b) to address the question whether the region is important for the symptom development and (2) AP_SAP11-like_PM19 N-terminally fused with a nuclear export signal (NES) of HIV-Rev (25), resulting in NES-AP_SAP11-like_PM19 (Fig. 1b), to analyze whether the localization of the protein has an effect on the development of symptoms. To prove the activity of the NES with our construct, we transiently expressed NES-AP_SAP11-like_PM19 fused to GFP in *N. benthamiana*. Microscopic analysis of the protoplast show that the protein is mainly localized in the cytoplasm (Fig. 2c).

The transgenic lines expressing AP_SAP11-like_PM19^Δ40-56^ showed a less severe phenotype compared to the plants expressing AP_SAP11-like_PM19, with a smaller rosette and increase in stem number (Fig. 3a) but the leaves and siliques resembling the wild type *A. thaliana* plant (Fig. 3b and c), suggesting that amino acids 40 to 56 of AP_SAP11-like_PM19 are somehow important for symptom development in *A. thaliana*, especially in the morphological changes of leaves and siliques. Similar results were reported for AY-WB_Sap11 (7).

Interestingly the transgenic *A. thaliana* lines expressing the NES-AP_SAP11-like_PM19 showed the same phenotype like the plant line expressing AP_SAP11-like_PM19 (Fig. 3a-c). Thus, the result indicates that the presence of AP_SAP11-like_PM19 only in cytoplasm is enough to induce this phenotype development.

To exclude the effect of the expression level of the transgene in the transgenic plant lines on the phenotype development, we performed RT-qPCR of homozygotes transgenic plant lines using an *AP_SAP11-like_PM19* specific primer that is able to bind to all three gene constructs, to ensure similar amplification. For each construct, three plant lines #1, #2 and #3 were analyzed. The transgenic plants shown in figure 3 a-c are represented by the respective line #1.

The maximal standard error of average Cq values of all three genes analyzed (*AP_SAP11-like_PM19* and reference genes *GAPDH* and *PP2A* (26) was 2.4 %, the maximal coefficient variation of Cq within the replicate samples was 7.1 %. The average efficiency of all three genes varied between 1.69 and 1.80. For a list of all received data see Table S2.

The obtained values were normalized to the arithmetic average of *AP_SAP11-like_PM19* for better comparison. The result in Fig. 3d shows that the relative expression level of AP_SAP11-like_PM19^Δ40-56^ and of NES_ AP_SAP11-like_PM19 are similar, although in average, they are slightly lower than the wild type AP_SAP11-like_PM19. This indicates that the different phenotypic developments are not caused by different expression levels. Since the transgenic plant expressing NES-AP_SAP11-like_PM19 still shows clear symptoms (Fig.3 a-c), we ruled out that the expression level is the cause of losing the crinkled siliques and leaves symptoms in the transgenic lines expressing the Δ40-56 mutant.

### Amino acids 40 to 56 of AP_SAP11-like_PM19are important for binding to some *A. thaliana* (At)TCPs in yeast two-hybrid (Y2H) analysis

Sugio and coworkers (6) showed that AY-WB_SAP11 interacted with class II AtTCP2 and 13 in Y2H screenings and with AtTCP2 and 4 of class II and AtTCP7 of class I in co-immunoprecipitation assays. Moreover a transient co-expression analysis showed that AY-WB_SAP11 destabilizes all class II AtTCP of the CIN-group, but not the class I AtTCP7 (6). Deletion of the N-terminal region of AY-WB_SAP11 including the NLS disturbs its ability to destabilize AtTCP-transcription factors in co-immunoprecipitation assay (7).

AP_SAP11-like protein of strain STAA interacts with three TCP transcription factors of *M. domestica* (MdTCP) that are homologous to members of AtTCP class II, namely MdTCP25 (a homolog of AtTCP4), MdTCP24, (a homolog of AtTCP13) and MdTCP16 (an isoform of AtTCP18) (10). *In planta* interaction of AP_SAP11-like protein strain STAA with MdTCP24 and 25 was confirmed using bimolecular fluorescence complementation (BiFC) (10). To our knowledge, so far, no interaction of AP_SAP11-like protein with members of AtTCP class I has been reported.

Since our results indicate that the amino acids 40 to 56 of AP_SAP11-like_PM19 are not responsible for nuclear localization of the protein, but responsible for some of the phenotypic characteristics of the transgenic *Arabidopsis* expressing AP_SAP11-like protein, therefore we analyzed its possible role during the interaction with AtTCP transcription factors using Y2H screens. In the first step, we screened for interaction of AP_SAP11-like_PM19 with all AtTCP transcription factors identified so far (27). We used the AP_SAP11-like protein fused to the Gal4 binding domain (BD) for bait (expression plasmid pGBKT7-*AP_SAP11-like_PM19*) and the AtTCP transcription factors fused to the Gal4 activation domain (AD) for prey (expression plasmid pGADT7-*AtTCP*). The pGBKT7-empty plasmid expressing only BD was included for negative control. Co-transformed yeast cells carrying the bait or pGBKT7 and one of the candidate prey plasmids were screened for resistance to Aureobasidin A (AbA) and expression of α-galactosidase. All Y2H results are given in supplementary material (Figure S1), examples for positive and negative Y2H results are shown in Figure 4. All Y2H results are summarized in Table1.

**Table 1:** Y2H results of AP_SAP11-like_PM19 and AP_SAP11-like_PM19^Δ40-56^ with different AtTCPs.

| AtTCP | Subclass | Interaction with AP_SAP11-like_PM19 | Interaction with AP_SAP11-like_PM19<sup>Δ40-56</sup> |
| --- | --- | --- | --- |
| 1 | Class II | pos. | pos. |
| 2 | Class II | pos. | pos. |
| 3 | Class II | pos. | neg. |
| 4 | Class II | pos. | neg. |
| 5 | Class II | pos. | pos. |
| 6 | Class I | pos. | pos. |
| 7 | Class I | pos. | pos. |
| 8 | Class I | neg. | n.a. |
| 9 | Class I | pos. | pos. |
| 10 | Class II | pos. | neg. |
| 11 | Class I | neg. | n.a. |
| 12 | Class II | pos. | pos. |
| 13 | Class II | pos. | pos. |
| 14 | Class I | pos. | pos. |
| 15 | Class I | neg. | n.a. |
| 16 | Class I | neg. | n.a. |
| 17 | Class II | pos. | pos. |
| 18 | Class II | pos. | pos. |
| 19 | Class I | pos. | neg. |
| 20 | Class I | neg. | n.a. |
| 21 | Class I | pos. | pos. |
| 22 | Class I | neg. | n.a. |
| 23 | Class I | neg. | n.a. |
| 24 | Class II | pos. | pos. |

pos. interaction of proteins in Y2H; neg. no interaction in Y2H; n.a. not applicable

**Figure 4:**
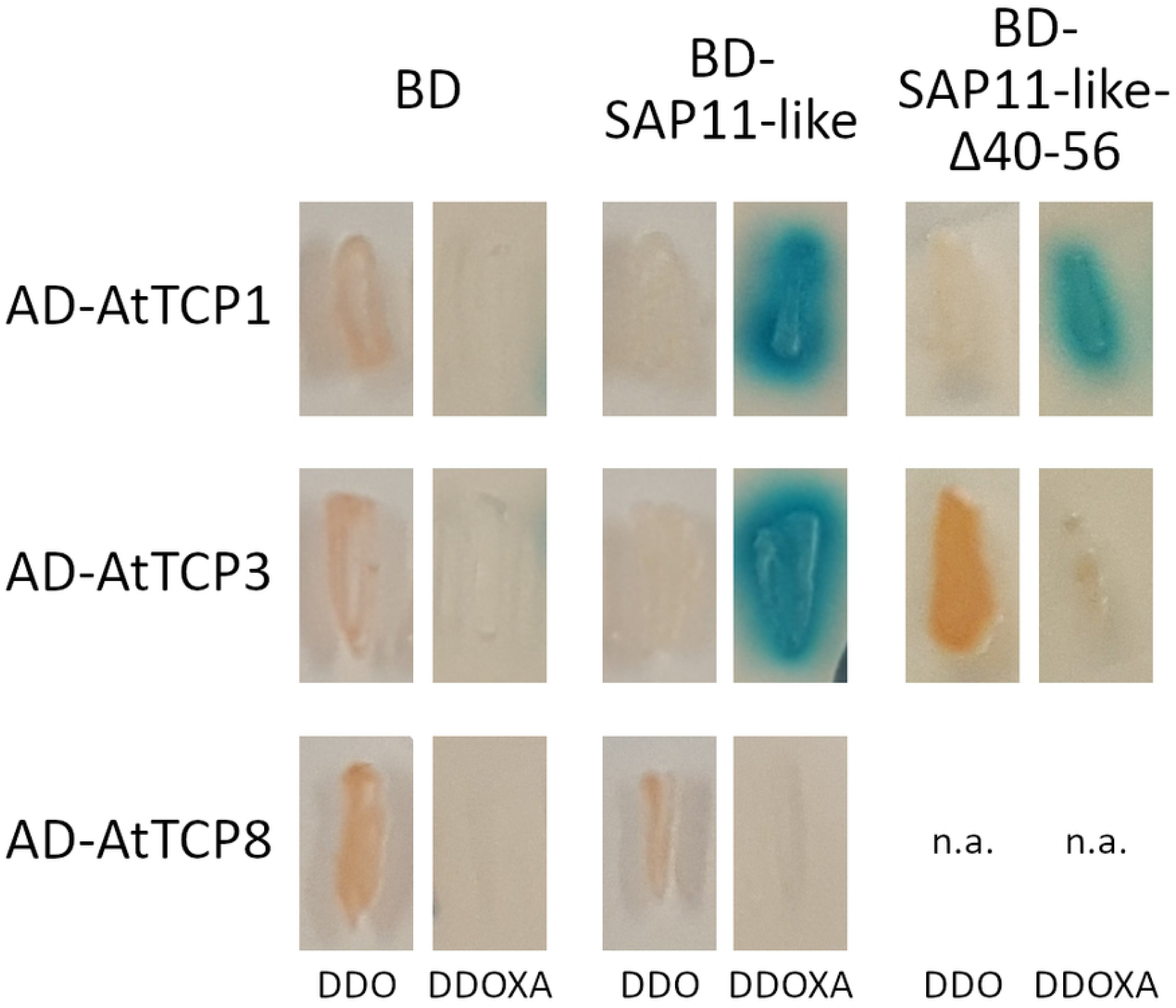
Example for Y2H results of AP_SAP11-like_PM19 and AP_SAP11-like_PM19^Δ40-56^ with different AtTCPs. Y2H screens were performed using the binding domain fused to AP_SAP11-like_PM19 (BD-SAP11-like) or AP_SAP11-like_PM19^Δ40-56^ (BD-SAP11-like-Δ40-56) and the activation domain (AD) fused to different AtTCPs. For negative control only BD was used. Cotransformed yeast cells were patched on double drop-out medium (DDO) to select for presence of both expression plasmids and DDO media containing Aureobasidin A and X-α-Gal (DDOXA) to select for protein interaction. n.a. = not applicable. A figure with all Y2H results can be found in supplementary material (Figure S1).

All eleven members of the AtTCPs class II (AtTCP1-5, 10, 12, 13, 17, 18 and 24) and six out of thirteen members of AtTCPs class I (AtTCP6, 7, 9, 14, 19 and 21) showed activation of both reporter genes in combination with AP_SAP11-like_PM19 whereas the remaining seven AtTCPs of class I (AtTCP8, 11, 15, 16, 20, 22 and 23) as well as all negative controls did not. This result indicates that AP_SAP11-like_PM19 protein does not only exclusively interact with class II but also with some of class I AtTCPs.

In the next step, we repeated Y2H screens of all candidate preys that showed interaction with AP_SAP11-like_PM19, using AP_SAP11-like_PM19^Δ40-56^ deletion mutant for bait. The result shows that AP_SAP11-like_PM19^Δ40-56^ binds to all tested AtTCPs, except AtTCP19 of class I and AtTCP3, 4 and 10 of class II. The results clearly indicate that the AP_SAP11-like_PM19^40-56^ stretch indeed is important for the interaction with some AtTCPs, especially some of the class II.

### Deletion of amino acids 40 to 56 of AP_SAP11-like_PM19 leads to a loss of its binding ability with AtTCP4 *in vivo* using FRET analysis

To prove the binding interaction results of the Y2H analysis *in vivo*, we performed FRET analysis using acceptor photo bleaching. This method is based on acceptor depletion and continuous monitoring of donor and acceptor fluorescence intensities during the acceptor photobleaching process (28). This method is reliable and has been wildly used to study protein interaction in the cells (29).

To analyze the interaction of the proteins in question in the plant cells, AP_SAP11-like_PM19 and AP_Sap11-like _PM19^Δ40-56^ proteins fused to GFP (donor) were transiently co-expressed with AtTCP4 (responding to leaf morphology, (30) or AtTCP13 fused to RFP (acceptor). The protoplasts were isolated and the binding interactions were investigated using acceptor FRET bleaching analysis. (Figure 5a-e). We observed that the intensity of the donor was increased after acceptor photobleaching for the FRET-pairs AP_SAP11-like _PM19-GFP/AtTCP4-RFP, AP_SAP11-like_PM19-GFP/AtTCP13-RFP and AP_SAP11-like_PM19^Δ40-56^/AtTCP13-RFP, indicating FRET had taken place while it was almost unchanged in AP_SAP11-like_PM19^Δ40-56^ /AtTCP4-RFP. The results were reproducible over a set of 10 protoplasts for each FRET pair.

**Figure 5:**
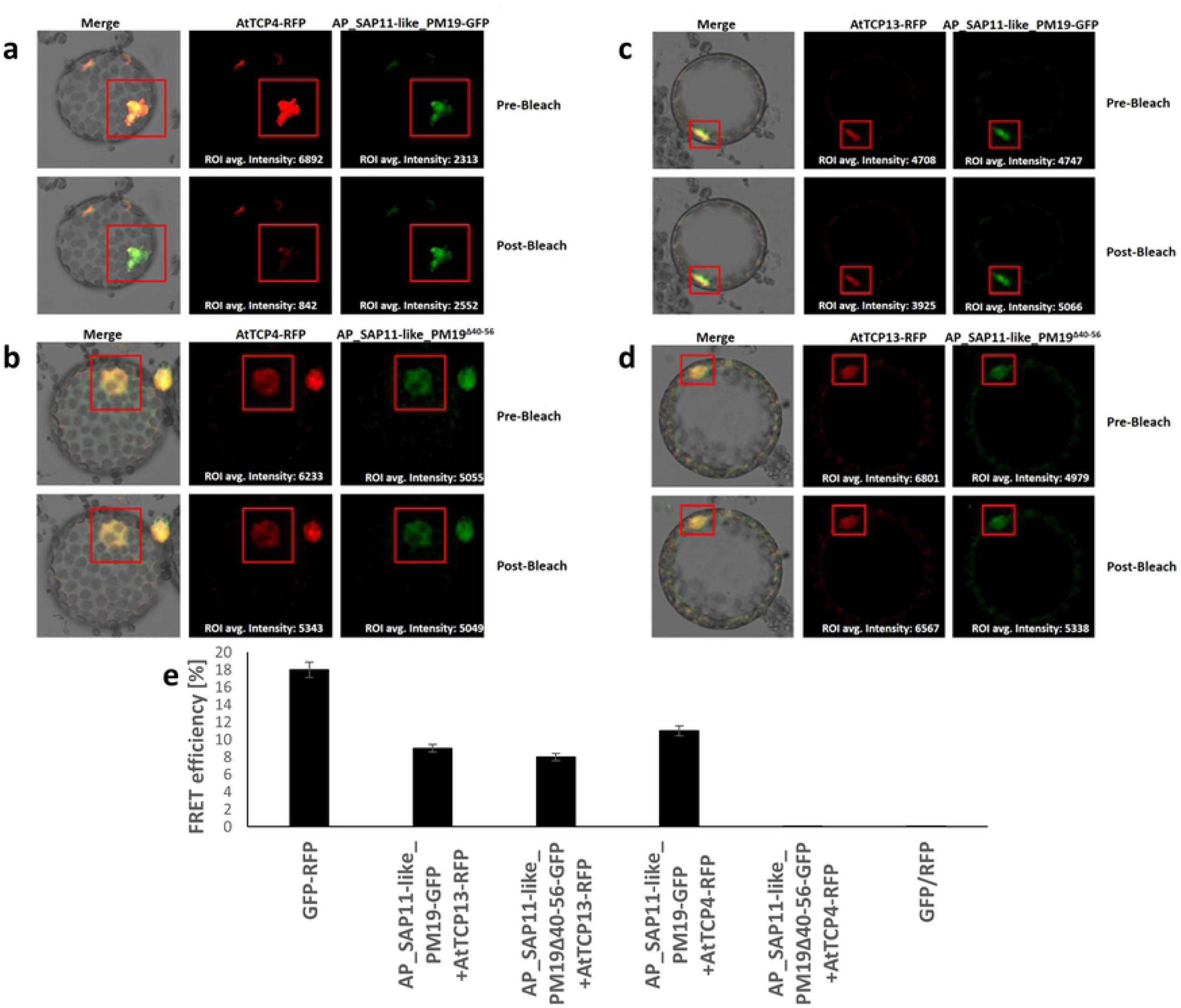
*In vivo* protein interaction assay using FRET analysis. AP_SAP11-like_PM19-GFP (**a.** and **c.**) and AP_SAP11-like _PM19^Δ40-56^ (**b.** and **d.**) were transiently expressed with AtTCP4 or AtTCP13-RFP and acceptor photobleaching FRET (abFRET) was performed. Both AP_SAP11-like_PM19-GFP and AP_SAP11-like_PM19^Δ40-56^ are mainly located in the nucleus. After abFRET, the RFP intensity is reduced while the GFP intensity increases in the protoplasts expressing AP_SAP11-like_PM19-GFP/AtTCP4-RFP (**a.**), AP_SAP11-like_PM19-GFP/AtTCP13-RFP (**c.**) and AP_SAP11-like_PM19^Δ40-56^ /AtTCP13-RFP (**d.**). The GFP signal almost not changed in the AP_SAP11-like_PM19^Δ40-56^/AtTCP4-RFP combination (**b.**). Squares indicate the region of interest (ROI) that was acceptor photobleached. **E.** Quantification of FRET photobleaching experiments by calculating FRET efficiencies for fusion protein RFP-GFP (positive control) is 17±2,3%, of AP_SAP11-like_PM19-GFP with AtTCP13-RFP it is 9±2.4 % while it is 8±2.02 % for AP_SAP11-like_PM19^Δ40-56^ with AtTCP13-RFP. FRET pair AP_SAP11-like_PM19-GFP/AtTCP4-RFP shows an efficiency of 11±2.4 % while it is 0.01 ±0.02 % for AP_SAP11-like_PM19^Δ40-56^ /AtTCP4-RFP. For positive FRET control protoplasts expressing RFP fused to GFP to obtain maximum FRET values were used, and for FRET negative control cells expressing RFP and GFP separately (from the same plasmid) to obtain the lowest FRET value (data not shown). Error bars indicate standard deviations. For all pairs, values of 10 protoplasts were used in the calculation % FRET efficiency (n = 10).

FRET efficiencies were calculated for all four FRET-pairs as well as a positive control (fusion protein GFP-RFP) and a negative control (separately expressed GFP and RFP). All FRET-pairs showed clearly higher FRET efficiencies than the separately expressed RFP and GFP (Figure 5e), except AP_SAP11-like_PM19^Δ40-56^/AtTCP4-RFP. The results clearly indicate that both AP_SAP11-like_PM19 and AP_SAP11-like_ PM19^Δ40-56^ bind AtTCP13, while the binding activity with AtTCP4 is lost for the Δ40-56-deletion mutant *in vivo*. Thus, the results clearly confirm the Y2H results *in vitro*.

However, in this experiment the resulting FRET-efficiencies of three pairs that indicated interaction of the acceptor and donor proteins were still clearly lower than the FRET-efficiency of the GFP-RFP fusion protein. *In vivo* AP_SAP11-like_PM19-GFP and AP_SAP11-like_PM19^Δ40-56^-GFP did not only interact with the transiently expressed AtTCP-RFP proteins, but also with endogenous AtTCPs of the plant cell. This leads to an underestimation of the FRET values compared to GFP-RFP as positive control.

## Discussion

The SAP11 effector protein of AY-WB phytoplasma is thought to play a major role in the infection mechanism of this phytoplasma. Its interaction with and destabilization of *A. thaliana* CIN-TCP transcription factors leads to a deep disturbance of plant biological processes resulting in morphological changes as well as a down-regulation of LOX2 expression and JA synthesis and an increase of insect vector reproduction (6). Although AP_SAP11-like protein bears little resemblance at amino acid level compared to AY-WP_SAP11, it was suggested that they may have some similar functions. Janik and co-workers proved interaction of AP_SAP11-like protein with some *M. domestica* TCP transcription factors and hence suggested an involvement in the hormonal changes occurring in apple trees during AP infection (10). In *N. benthamiana*, AP_SAP11-like protein destabilizes TCP transcription factors, supress JA response and alter the aroma phenotype, that might play a role in the attraction of the insect vector, as well as in the morphology of transgenic plants (13).

Despite their similar effect on plant development and plant hormone biosynthesis, so far, no protein domains have been identified that match each other in both proteins (AP_SAP11-like protein and AY-WB_SAP11), leaving functions of different areas of the AP_SAP11-like protein on a hypothetical basis. In this work, we focus on elucidating the role of the amino acid 40-56 stretch of AP_SAP11-like_PM19 protein on the localization of the protein in plant cells, the phenotype development in transgenic *A. thaliana* lines and on the AtTCP-binding activity.

We show that AP_SAP11-like_PM19 protein is, like AY-WB_SAP11, mainly located in the plant cell nucleus when fused to GFP and transiently expressed in *N. benthamiana*. But the amino acids 40 to 56 corresponding to the AY-WB NLS in sequence alignment, do not responding for the nuclear import of this protein. The same finding was reported for a SAP11 like effector, SWP1 isolated from wheat blue dwarf phytoplasma. It was shown that deletion of the C-terminal-predicted NLS of the protein did not affect the localization of the protein into the nucleus (14).

Fusing a small protein with GFP for nuclear localization study in plant cells could result in miss interpretation, as GFP can enter to the nucleus through nuclear pores (23). Moreover small proteins up to 60 kDa can passively enter the nucleus (22). Since AP_SAP11-like_PM19 is a small protein (11 kDa) and our results indicate that it does not contain an actual NLS as well as it fails to locate into the nucleus when its size is increased (fused with Gus-GFP), it cannot be ruled out that the protein could enter the nucleus via passive transport or other unknown mechanisms.

It was reported that NLS of AY-WB_SAP11 protein is not required for binding some AtTCPs (7). However, deletion of the N-terminal area of AY-WB_SAP11 including the NLS disrupts its ability to destabilize some AtTCPs which leads to mild symptoms in *A. thaliana* transgenic plant lines (7).To investigate the role of the amino acid 40-56 stretch of AP_SAP11-like_PM19 during symptom development, different transgenic plant lines expressing AP_SAP11-like_PM19, a mutant missing amino acids 40 to 56, or a fusion with nuclear export signal (NES) were generated. The results show that the *A. thaliana* lines expressing AP_SAP11-like_PM19 and NES-fused AP_SAP11-like_PM19 proteins show the same typical phenotypes (crinkled leaves and siliques and witches’ broom) as were reported for plants expressing AY-WB_SAP11 (6). However, the plants expressing AP_SAP11-like_PM19^Δ40-56^ show no crinkled leaves and siliques, but the witches’ broom phenotype remains. This is consistent with the findings for AY-WB phytoplasma (6, 14). The RT-PCR results support the finding that the partial loss of the crinkled leaves and siliques phenotype in the plants expressing AP_SAP11-like_PM19^Δ40-56^ is not due to the expression level of the transgene.

It was discussed that the AY-WB_SAP11 enters the nucleus to interact with and destabilize AtTCPs, however a cytoplasmic function of the protein was not completely excluded (7). Since our protein localization results show that AP_SAP11-like_PM19 and the Δ40-56 mutant are distributed in the plant cell in the same manner, the effect of amino acids 40-56 on the protein localization can be excluded. Moreover the transgenic plant lines expressing NES-AP_SAP11-like_PM19 where the expressed protein is exclusively localized in the cytoplasm, develop the same phenotypes as the plants expressing AP-SAP11-like_PM19, supporting the assumption that the nuclear localization of the expressed protein is not the major cause for losing phenotypes in the transgenic plants expressing AP_SAP11-like_PM19^Δ40-56^.

The result of the Y2Hscreening, the interaction of AP_SAP11-like_PM19 and the Δ40-56 mutant with AtTCPs, indicate that the AP_SAP11-like_PM19^Δ40-56^-stretch is not required for binding all AtTCPs but it is necessary for binding of AtTCPs 3, 4, 10 and 19. The requirement of the amino acids 40-56-stretch for binding with at least AtTCP4 was confirmed *in vivo* using acceptor photobleaching FRET analysis.

TCP transcription factors have been intensively studied in the past years, showing that they regulate a variety of plant processes from plant development to defence responses. The functions of different AtTCPs and their role in biosynthetic processes have been reviewed in detail by Shutian Li (31). The variety of their functions reaches from developmental processes over involvement in the clock oscillator to defence response (31).

We showed that the amino acids 40-56 stretch of AP_SAP11-like_PM19 is important for binding to class II AtTCPs 3, 4 and 10. These three AtTCPs are closely related to one another (31) and belong to a group of five AtTCPs that are regulated by the microRNA miR319 (30). Overexpression of miR319 leads to down regulation of AtTCPs 2, 3, 4, 10 and 24 and a crinkled leaf phenotype (30). Crinkled leaves and/or siliques have also been observed in other *A. thaliana* mutants when the members of this group of AtTCPs are down regulated (32, 33). These findings together with our Y2H screening results and FRET analysis with AtTCP4 strongly suggest that the loss of binding activity with AtTCP 3, 4 and 10 of AP_SAP11-like_PM19^Δ40-56^ results in disappearance of the crinkled leaves and siliques phenotype in the transgenic plants.

However, the transgenic *A. thaliana* lines expressing AP_SAP11-like_PM19^Δ40-56^ still shows witches’ broom phenotype and smaller rosettes. This could be due to the bindings of AP_SAP11-like_PM19^Δ40-56^ to AtTCP12 and 18, which redundantly control branch overgrowth (34–36). The same result was obtained in transgenic plant expressing the deletion of predicted NLS of SAP11 like effector, SWP1 (14).

## Conclusion

Taking all results into account, the results strongly suggest that the amino acids 40-56 -stretch of AP_SAP11-like_PM19 is not important for nuclear localization of the protein and is at least a part of the binding site with AtTCPs 3, 4, 10 and 19 which are important in controlling the plant leaf morphology. Moreover, the AP_SAP11-like_PM19 can not only bind all AtTCPs of class II but also some of class I.

## Material and Methods

### Origin of AP_SAP11-like_PM19 DNA

‘*Ca*. P. mali’ strain PM19 was previously transmitted from field-collected *Cacopsylla picta* to healthy test plants of *Malus x domestica* (15). Total DNA was extracted from plant tissue using a modified cetyltrimethylammonium bromide (CTAB)-based protocol described elsewhere (15). The gene *AP_SAP11-like_PM19* (GenBank Accession number MK966431) without SMV signal was amplified using primer that were designed based on its corresponding gene of ‘*Ca*. P. mali’ strain AT (16): 5’-TCTCCTCCTAAAAAAGATTC-3’ and 5’-TTTTTTTCCTTTGTCTTTATTGTT-3’.

### Transient protein expression *in planta* and protoplast isolation

The *AP_SAP11-like_PM19* genes was codon optimized for expression in *A. thaliana* and synthesized (GeneCust, Ellange, Luxembourg). This version of the gene was used as basis for all constructs used in *in planta* experiments in this work.

Agroinfiltration was performed as previously described (37). A single colony of *Agrobacterium tumefaciens* (*A. tumefaciens*) strain ATHV transformed with pPZP200 binary vector (38) containing the respective gene under control of the *Cauliflower mosaic virus* 35S promoter was grown at 28 °C. Bacteria were centrifuged, resuspended in induction media (bacterial growth medium substituted with 10 mM MES, 2 mM MgSO_4_ and 0.05 M acetosyringone) and grown over night at 28 °C. After centrifuging, the cell pellet was resuspended to an OD_600_ of 2.4 in infection media (1/2 MS media, 10 mM MES, 2% sucrose and 0.2 mM acetosyringone) and incubated at room temperature for at least 2 h. The bacteria were infiltrated into *N. benthamiana* leaves using a 1 mL needleless syringe.

For protoplast isolation, two leaves were collected 2 days after infiltration, cut in small strips of about 1 mm and placed in enzyme solution for cell wall maceration [1/2 MS, 0.4 M sucrose, 1% Cellulase Onozuka R-10 and 0.2 % Macerozyme R-10 (both Serva, Heidelberg, Germany), pH = 5.8]. After infiltration by application of vacuum, leaves were incubated for 2 hours on a gyratory shaker (20 rpm). Protoplasts were collected by centrifuging at 100 g and examined using a Zeiss Observer Z1 with LSM510 confocal laser-scanning head.

### Generation of transgenic *A. thaliana* lines

Floral dip was performed as previously described (39). A single colony of *A. tumefaciens* strain GV3101 transformed with pPZP200 binary vector (38) containing the respective gene under control of the *Cauliflower mosaic virus* 35S promoter was grown over night at 28 °C. Bacteria were centrifuged, resuspended in infection media (1/2 MS, 10 mM MES; 5 % sucrose, 0.2 mM acetosyringone and 0.05 % Silwet L-77) and incubated at room temperature for at least 2 hours. Flowers of *A. thaliana* plants were submerged in the bacteria suspension for 30 seconds. Plants were kept in a dark and humid area for 2 days and then grown in greenhouse in long-day (16 h/8 h light/dark) conditions until seeds could be collected. F1 and F2 generation of transgenic plants were selected by spraying of BASTA solution, diluted 1/1000 in H_2_O. The F2 generation of transgenic *A. thaliana* lines was screened for phenotypic symptoms and analyzed using RT-qPCR.

### RT-qPCR

For RT-qPCR, RNA of transgenic *A. thaliana* plants was extracted using the Macherey-Nagel RNA extraction kit (Macherey-Nagel, Düren, Germany). Resulting RNA was additionally treated with DNase I in solution (according to kit protocol) and tested for DNA contamination by performing RT-qPCR with 10 ng and 50 ng RNA per reaction. Only samples that did not show any signal in both reactions were used for cDNA synthesis. cDNA was synthesized from 0.5 µg of total RNA using a RevertAid Premium Kit (Fermentas) and subjected to RT-qPCR using iTaq Universal SYBR Green Supermix (BioRad) and 10 ng of cDNA per reaction. RT-qPCR reaction was performed in a Chromo 4™ system cycler (Biorad). Four technical replicates were produced per plant by repeating the cDNA synthesis step separately twice and performing two RT-qPCR reactions per cDNA sample. Two reference genes were used: *glyceraldehyde-3-phosphate dehydrogenase* (*GAPDH*) and *protein phosphatase 2* (*PP2A*). *GAPDH* is a standard reference gene, often used in RT-qPCR (26). *PP2A* is one of many genes, suggested as new reference genes by Czechowski and coworkers due to their stable expression (26). For *GAPDH* and *PP2A* primers were used as described (26). For amplification of *AP_SAP11-like_PM19* and its mutations, gene-specific primers that can bind to all three variants, were designed (supplementary material, Table S1). The mean Cq values of the technical replicates, as well as standard error of average Cq values, the maximal coefficient variation of Cq within replicate samples and the average efficiency of genes were calculated using the Real-Time PCR Miner software (40) and are stated in supplementary material, Table S2 and S3 respectively.

The relative expression levels were calculated using the ddCt method (41) and normalized to the geometric average of the Cq of the reference genes. In the calculation, the relative expression level of each gene was also normalized to that of *AP_SAP11-like_PM19*. The exact equation used for calculation is stated in equation 1. 

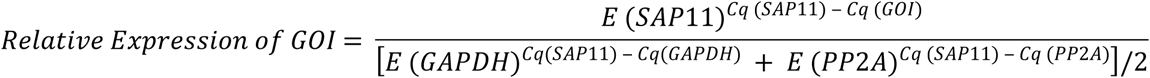

**Equation 1: Equation used for calculation of relative expression levels of gene of interest (GOI)**. Average Cq values of GOI were normalized to the geometric average of the Cq values of *glyceraldehyde-3-phosphate dehydrogenase* (*GAPDH*) and *protein phosphatase 2* (*PP2A*) and to the relative expression level of *AP_SAP11-like_PM19*. E is average efficiency of gene amplification.

### Yeast two-hybrid analysis

Y2H screening was performed using Matchmaker Gold Yeast Two-Hybrid System (Takara Bio USA, Inc., Mountain View, US-CA). For bait *AP_SAP11-like_PM19* or *AP_SAP11-like_PM19*^*Δ40-56*^ was cloned into pGBKT7 (TRP1 nutritional marker) for fusion with the Gal4 binding domain (*BD*). Empty pGBKT7 vector expressing only BD was used for negative control. For prey, DNA samples with the genes of all 24 AtTCP transcription factors were purchased from The Arabidopsis Information Resource (TAIR, Phoenix Bioinformatics, Fremont, USA-CA), PCR amplified and cloned into pGADT7 (LEU2 nutritional marker) for fusion with the Gal3 activation domain (*AD*). Different yeast drop-out media were purchased from Sigma-Aldrich Inc. (St. Louis, US-MO) and prepared according to manufacturer’s instructions. 40 µg/mL X-α-Gal (Iris Biotech GmbH, Marktredwitz, Germany) and 200 ng/mL AbA (Takara Bio USA, Inc., Mountain View, US-CA) were added when they are required. Two reporter genes were used to detect protein interaction: AUR1-C confers strong resistance to Aureobasidin A (Aba); MEL1 encodes α-Galactosidase. When the substrate X-α-Gal is added to the media, positive colonies turn blue.

To test for interaction between bait and prey, the respective expression plasmids were co-transformed into yeast strain Y2H gold and patched on ^−^Leu/Trp double drop out plates (DDO) that selects for presence of both expression plasmids and DDO plates substituted with X-α-Gal and Aba (DDOXA) to select for positive interactions.

### Acceptor photobleaching FRET

The genes of the two potential interaction partners fused to *GFP* (donor) and *RFP* (acceptor) respectively, were cloned into the same pPZP200 binary vector (38), to ensure coexpression in the same plant cell. The proteins were then transiently expressed in *N. benthamiana* and protoplasts were isolated, as described before.

For fluorescence resonance energy transfer (FRET) analysis the signal of the donor GFP was observed using a Zeiss Observer Z1 with LSM510 confocal laser-scanning head before and after photobleaching of the acceptor RFP. An increase of GFP signal indicates protein-protein-interaction of the two potential partners. The FRET efficiency was calculated with equation E = 1-F_DA_/F_D_ where F_DA_ is the fluorescence intensity of the donor in the presence of the acceptor (pre-bleach) and F_D_ is the fluorescence intensity of the donor when the acceptor is far away (post-bleach) (42).

## Acknowledgement

We thank Dr. Wolfgang Jarausch for providing DNA extract from *‘Ca.* P. mali’ strain PM19.

## Supplementary information

**Figure S1: Y2H results of AP_SAP11-like_PM19 and AP_SAP11-like_PM19**^**Δ40-56**^ **with different AtTCPs.** Y2H screens were performed using the binding domain fused to AP_SAP11-like_PM19 (BD-SAP11-like) or AP_SAP11-like_PM19^Δ40-56^ (BD-SAP11-like-Δ40-56) and the activation domain (AD) fused to different AtTCPs. For negative control only BD was used. All negative controls showed expected results (left column). AP_SAP11-like_PM19 binds to six AtTCP of class I and all of classII (middle column). Binding ability with AtTCP3, 4, 10 and 19 is lost, when aa 40 to 56 are deleted, while it remains with the other AtTCPs (right column). Results of these Y2H experiments are also summarized in Table 1.

**Table S1: Gene-specific primer used for RT-qPCR.**

**Table S2: Average Cq values, standard error and maximal coefficient variation of Cq values within replicate samples of RT-qPCR**

## Literature Cited

1. Hogenhout SA, van der Hoorn RAL, Terauchi R, Kamoun S. Emerging concepts in effector biology of plant-associated organisms. Mol Plant Microbe Interact 2009; 22(2):115–22.

2. Sugio A, MacLean AM, Kingdom HN, Grieve VM, Manimekalai R, Hogenhout SA. Diverse targets of phytoplasma effectors: from plant development to defense against insects. Annu Rev Phytopathol 2011; 49:175–95.

3. Hogenhout SA, Oshima K, Ammar E-D, Kakizawa S, Kingdom HN, Namba S. Phytoplasmas: bacteria that manipulate plants and insects. Mol Plant Pathol 2008; 9(4):403–23.

4. Bai X, Zhang J, Ewing A, Miller SA, Jancso Radek A, Shevchenko DV et al. Living with genome instability: the adaptation of phytoplasmas to diverse environments of their insect and plant hosts. J Bacteriol 2006; 188(10):3682–96.

5. Bai X, Correa VR, Toruño TY, Ammar E-D, Kamoun S, Hogenhout SA. AY-WB phytoplasma secretes a protein that targets plant cell nuclei. Mol Plant Microbe Interact 2009; 22(1):18–30.

6. Sugio A, Kingdom HN, MacLean AM, Grieve VM, Hogenhout SA. Phytoplasma protein effector SAP11 enhances insect vector reproduction by manipulating plant development and defense hormone biosynthesis. Proc Natl Acad Sci U S A 2011; 108(48):E1254–63.

7. Sugio A, MacLean AM, Hogenhout SA. The small phytoplasma virulence effector SAP11 contains distinct domains required for nuclear targeting and CIN-TCP binding and destabilization. New Phytol 2014; 202(3):838–48.

8. Siewert C, Luge T, Duduk B, Seemüller E, Büttner C, Sauer S et al. Analysis of expressed genes of the bacterium ‘Candidatus phytoplasma Mali’ highlights key features of virulence and metabolism. PLoS ONE 2014; 9(4):e94391.

9. Jomantiene R, Zhao Y, Davis RE. Sequence-variable mosaics: composites of recurrent transposition characterizing the genomes of phylogenetically diverse phytoplasmas. DNA Cell Biol 2007; 26(8):557–64.

10. Janik K, Mithöfer A, Raffeiner M, Stellmach H, Hause B, Schlink K. An effector of apple proliferation phytoplasma targets TCP transcription factors-a generalized virulence strategy of phytoplasma? Mol Plant Pathol 2017; 18(3):435–42.

11. Mayer CJ, Vilcinskas A, Gross J. Pathogen-induced release of plant allomone manipulates vector insect behavior. J Chem Ecol 2008; 34(12):1518–22.

12. Mayer CJ, Vilcinskas A, Gross J. Phytopathogen lures its insect vector by altering host plant odor. J Chem Ecol 2008; 34(8):1045–9.

13. Tan CM, Li C-H, Tsao N-W, Su L-W, Lu Y-T, Chang SH et al. Phytoplasma SAP11 alters 3-isobutyl-2-methoxypyrazine biosynthesis in Nicotiana benthamiana by suppressing NbOMT1. J Exp Bot 2016; 67(14):4415–25.

14. Wang N, Yang H, Yin Z, Liu W, Sun L, Wu Y. Phytoplasma effector SWP1 induces witches’ broom symptom by destabilizing the TCP transcription factor BRANCHED1. Mol Plant Pathol 2018; 19(12):2623–34.

15. Jarausch B, Schwind N, Fuchs A, Jarausch W. Characteristics of the spread of apple proliferation by its vector Cacopsylla picta. Phytopathology 2011; 101(12):1471–80.

16. Kube M, Schneider B, Kuhl H, Dandekar T, Heitmann K, Migdoll AM et al. The linear chromosome of the plant-pathogenic mycoplasma ‘Candidatus Phytoplasma mali’. BMC Genomics 2008; 9:306.

17. Beckwith J. The Sec-dependent pathway. Res Microbiol 2013; 164(6):497–504.

18. Wada Y, Ohya H, Yamaguchi Y, Koizumi N, Sano H. Preferential de novo methylation of cytosine residues in non-CpG sequences by a domains rearranged DNA methyltransferase from tobacco plants. J Biol Chem 2003; 278(43):42386–93.

19. Brameier M, Krings A, MacCallum RM. NucPred--predicting nuclear localization of proteins. Bioinformatics 2007; 23(9):1159–60.

20. Cokol M, Nair R, Rost B. Finding nuclear localization signals. EMBO Rep 2000; 1(5):411–5.

21. Nakai K, Horton P. PSORT: a program for detecting sorting signals in proteins and predicting their subcellular localization. Trends Biochem Sci 1999; 24(1):34–6.

22. Wang R, Brattain MG. The maximal size of protein to diffuse through the nuclear pore is larger than 60kDa. FEBS Lett 2007; 581(17):3164–70.

23. Seibel NM, Eljouni J, Nalaskowski MM, Hampe W. Nuclear localization of enhanced green fluorescent protein homomultimers. Anal Biochem 2007; 368(1):95–9.

24. Grebenok RJ, Pierson E, Lambert GM, Gong FC, Afonso CL, Haldeman-Cahill R et al. Green-fluorescent protein fusions for efficient characterization of nuclear targeting. Plant J 1997; 11(3):573–86.

25. Fischer U, Huber J, Boelens WC, Mattaj IW, Lührmann R. The HIV-1 Rev activation domain is a nuclear export signal that accesses an export pathway used by specific cellular RNAs. Cell 1995; 82(3):475–83.

26. Czechowski T, Stitt M, Altmann T, Udvardi MK, Scheible W-R. Genome-wide identification and testing of superior reference genes for transcript normalization in Arabidopsis. Plant Physiol 2005; 139(1):5–17.

27. Danisman S, van der Wal F, Dhondt S, Waites R, Folter S de, Bimbo A et al. Arabidopsis class I and class II TCP transcription factors regulate jasmonic acid metabolism and leaf development antagonistically. Plant Physiol 2012; 159(4):1511–23.

28. Karpova TS, Baumann CT, He L, Wu X, Grammer A, Lipsky P et al. Fluorescence resonance energy transfer from cyan to yellow fluorescent protein detected by acceptor photobleaching using confocal microscopy and a single laser. J Microsc 2003; 209(Pt 1):56–70.

29. Skruzny, Pohl, Abella. FRET Microscopy in Yeast. Biosensors 2019; 9(4):122.

30. Schommer C, Palatnik JF, Aggarwal P, Chételat A, Cubas P, Farmer EE et al. Control of jasmonate biosynthesis and senescence by miR319 targets. PLoS Biol 2008; 6(9):e230.

31. Li S. The Arabidopsis thaliana TCP transcription factors: A broadening horizon beyond development. Plant Signal Behav 2015; 10(7):e1044192.

32. Qin G, Gu H, Zhao Y, Ma Z, Shi G, Yang Y et al. An indole-3-acetic acid carboxyl methyltransferase regulates Arabidopsis leaf development. Plant Cell 2005; 17(10):2693–704.

33. Palatnik JF, Allen E, Wu X, Schommer C, Schwab R, Carrington JC et al. Control of leaf morphogenesis by microRNAs. Nature 2003; 425(6955):257–63.

34. Aguilar-Martínez JA, Poza-Carrión C, Cubas P. Arabidopsis BRANCHED1 acts as an integrator of branching signals within axillary buds. Plant Cell 2007; 19(2):458–72.

35. Poza-Carrión C, Aguilar-Martínez JA, Cubas P. Role of TCP Gene BRANCHED1 in the Control of Shoot Branching in Arabidopsis. Plant Signal Behav 2007; 2(6):551–2.

36. Finlayson SA. Arabidopsis Teosinte Branched1-like 1 regulates axillary bud outgrowth and is homologous to monocot Teosinte Branched1. Plant Cell Physiol 2007; 48(5):667–77.

37. Schöb H, Kunz C, Meins F. Silencing of transgenes introduced into leaves by agroinfiltration: a simple, rapid method for investigating sequence requirements for gene silencing. Mol Gen Genet 1997; 256(5):581–5.

38. Hajdukiewicz P, Svab Z, Maliga P. The small, versatile pPZP family of Agrobacterium binary vectors for plant transformation. Plant Mol Biol 1994; 25(6):989–94.

39. Clough SJ, Bent AF. Floral dip: a simplified method for Agrobacterium-mediated transformation of Arabidopsis thaliana. Plant J 1998; 16(6):735–43.

40. Zhao S, Fernald RD. Comprehensive algorithm for quantitative real-time polymerase chain reaction. J Comput Biol 2005; 12(8):1047–64.

41. Pfaffl MW. A new mathematical model for relative quantification in real-time RT-PCR. Nucleic Acids Res 2001; 29(9):e45.

42. Lakowicz JR. Principles of fluorescence spectroscopy [pp. 367-371]. Third edition, corrected at 4. printing. New York, NY: Springer; 2010.

